# Axo-axonic synaptic input drives homeostatic plasticity by tuning the axon initial segment structurally and functionally

**DOI:** 10.1101/2024.04.11.589005

**Authors:** Rui Zhao, Baihui Ren, Yujie Xiao, Jifeng Tian, Yi Zou, Jiafan Wei, Yanqing Qi, Ankang Hu, Xiaoying Xie, Z. Josh Huang, Yousheng Shu, Miao He, Jiangteng Lu, Yilin Tai

## Abstract

The stability of functional brain network is maintained by homeostatic plasticity, which restores equilibrium following perturbation. As the initiation site of action potentials, the axon initial segment (AIS) of glutamatergic projection neurons (PyNs) undergoes dynamic adjustment that exerts powerful control over neuronal firing properties in response to changes in network states. Although AIS plasticity has been reported to be coupled with the changes of network activity, it is poorly understood whether it involves direct synaptic input to the AIS. Here we show that changes of GABAergic synaptic input to the AIS of cortical PyNs, specifically from chandelier cells (ChCs), are sufficient to drive homeostatic tuning of the AIS within 1-2 weeks, while those from parvalbumin-positive basket cells do not. This tuning is reflected in the morphology of the AIS, the expression level of voltage-gated sodium channels, and the intrinsic neuronal excitability of PyNs. Interestingly, the timing of AIS tuning in PyNs of the prefrontal cortex corresponds to the recovery of changes in social behavior caused by alterations of ChC synaptic transmission. Thus, homeostatic plasticity of the AIS at postsynaptic PyNs may counteract deficits elicited by imbalanced ChC presynaptic input.

**Teaser:** Axon initial segment dynamically responds to changes in local input from chandelier cells to prevent abnormal neuronal functions.

## Introduction

The axon initial segment (AIS) is a highly specialized cellular compartment localized at the proximal axon of the neuron. It serves as a gatekeeper by filtering the somato-dendritic proteins to maintain the neuronal polarity (*1–4*). More importantly, AIS contributes to the initiation of action potential (AP) as it provides docking sites for high density of voltage-gated sodium channels, which makes AIS the most excitable part of a neuron (*5–9*). The precise position relative to the cell body, the morphology of the AIS, the voltage-gated ion channel expression at the AIS, as well as axo-axonic synaptic input to the AIS could have significant impacts on AP generation, propagation, and temporal precision (*10–16*). Therefore, subtle changes in the AIS structure or molecular composition are capable to regulate the neuronal output.

Neurons have the ability to tune themselves to maintain the stability of the network at multiple levels (*17*). Recent studies have shown that AIS undergoes plastic changes in response to alterations in network activity (*11, 12, 18, 19*). The length of the AIS and its precise location relative to the cell body dynamically adapt to global depolarization in cultured dentate granule cells (*11*). Additionally, sensory deprivation by either removing cochlea or depriving the whisker-to-barrel pathway leads to the elongation of the AIS, coupled with changes in ion channel expression and increased intrinsic excitability (*12, 19*). Conversely, increased network activity through sensory input or exposure to an enriched environment induces the shortening of the AIS and shapes the channel distribution and neuronal intrinsic excitability (*19, 20*). Such homeostatic adaptations of AIS at the structural and functional levels are believed to contribute to regulation of neuronal output (*21*), counteracting pathological conditions under injury or disease, thereby maintaining the stability of dynamic network activity (*22*).

While chronic (hours to days) alteration of network activity can result in homeostatic scaling of the AIS at the postsynaptic pyramidal neuron (PyN), another form of regulation exerts direct impact on neuronal output from the AIS: the axo-axonic synaptic input to the AIS from the axo-axonic cells (AACs). As a unique subdomain on the axon that receives synaptic input, the AIS of neocortical PyN receives synaptic inputs predominantly from a distinctive type of AACs, the chandelier cells (ChCs) (*23, 24*). By positioning their axonal terminals (called “cartridges”) on the AIS to form GABAergic axo-axonic synapses, ChCs are thought to exert powerful control over PyN firing to suppress irrelevant and excessive excitatory activity in neuronal networks (*25–29*). In the basal lateral amygdala (BLA), a single AP from a single AAC is sufficient to initiate sharp wave ripples (SWRs) through the local circuit (*30*). The strong impact of AACs on PyN firing raises the question of whether local alterations of AACs input are sufficient to drive PyN homeostatic plasticity by tuning the AIS and whether it contributes to functional adaptation at the cellular and behavioral levels.

In this study, we investigated the impact of chronic alterations of axo-axonic synaptic input on the AIS structure and function, as well as on the behavior of mice. Ablation of ChCs resulted in the shortening of the AIS and increased threshold of AP generation, coupled with decreased expression of voltage-gated sodium channels. Conversely, chronic activation of ChCs had the opposite effect. Interestingly, changes in social behavior caused by alterations of axo-axonic synaptic transmission were restored when AIS scaling had happened. In summary, our results indicate that prolonged changes of presynaptic ChCs inputs lead to modulations of postsynaptic structure and molecular changes at the AIS of PyNs, which serves to counteract the disturbed micro-circuitry at the functional level.

## Results

### Genetic ablation of ChCs leads to reduced synaptic input to the AIS

To assess the impact of direct synaptic input on AIS plasticity, we used an intersectional genetic strategy to label and manipulate ChCs (*31*). To do this, two sets of driver lines are involved (Figure 1A). The first driver line was the *knock-in* mouse driver line, Unc5b-2A-Cre^ERT2^ (referred to as *Unc5b-CreER* below)(*25*), which was created based on the fact that Unc5b expression is largely restricted to ChCs (*32*). However, crossing *Unc5b-CreER* driver line with the *Ai14* reporter line (*Rosa26-loxpSTOPloxp-tdTomato*) resulted in tdTomato expression in both ChCs and capillary endothelial cells (Figure S1). Therefore, we introduced the second driver line, the *Nkx2.1-Flp*, in which Flippase (Flp) is driven from the Nkx2.1 promotor. The Nkx2.1 lineage includes most of the interneurons born from medial ganglion eminence (MGE), of which the ventral area is where ChCs are born (*33, 34*). By combining the *Unc5b-CreER* and *Nkx2.1-Flp* driver lines with the dual recombinase-dependent reporter line, *Ai65,* we were able to specifically label ChCs with high efficiency (Figure S2A and S2B). In the dorsal medium prefrontal cortex (dmPFC), genetically labeled neurons were particularly enriched in cortical layer 2 (L2) with the typical chandelier-like morphology. The axonal terminals innervated the AIS of the PyNs, forming the “cartridges” with multiple boutons (Figure S2B1-B5). No capillary endothelial cells were observed, indicating that the intersectional strategy allowed for restricted gene expression in ChCs.

**Fig. 1.**
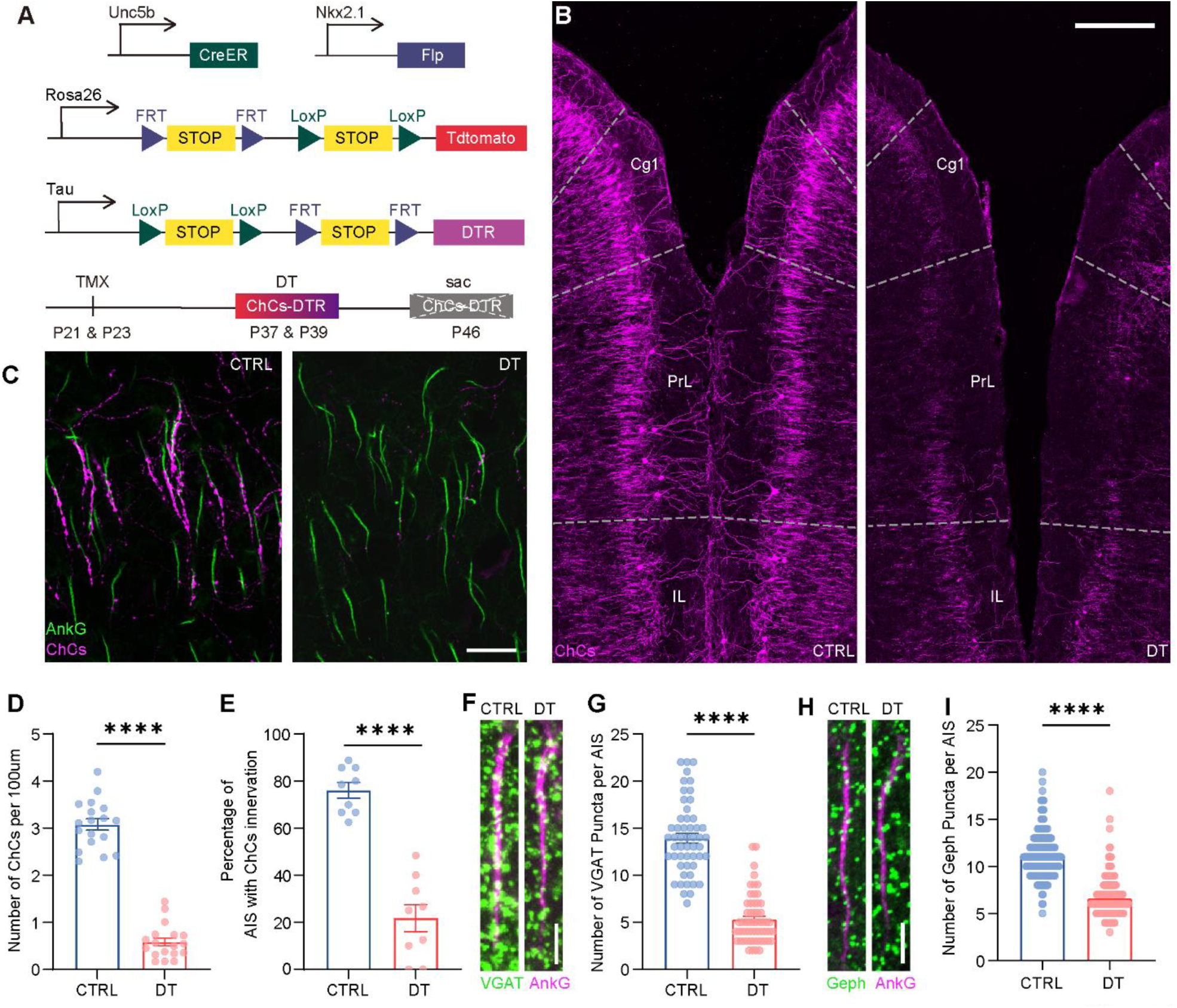
Strategy to genetically ablate ChCs in vivo. (**A**) Schematic representation of the strategy of intersectional labeling (tdTomato) and ablation (DTR: diphtheria toxin receptor) of the ChCs. TMX was administered at P21 and P23 to induce DTR expression, following by i.p. injection of DT (diphtheria toxin) at P37 and P39. Animals were sacrificed one week later to assess ablation efficiency. **(B)** Representative images showing successful ablation of ChCs in the dmPFC one week after DT injection. Scale bar: 200 μm **(C)** Representative images showing reduced innervation of the AISs by ChCs. Scale bar: 50 μm **(D)** Reduced number of ChCs per 100 μm located at the boundary of layer1 and layer 2 after genetic ablation (CTRL: *n* = 18 brain slices, 3 mice; DT: *n* = 18 brain slices, 3 mice; *P*<0.0001, unpaired *t*-test). **(E)** Decreased overall percentage of layer2/3 AISs innervated by ChC cartridges in the DT-treated group compared to the control group (CTRL: *n* = 9 brain slices, 3 mice; DT: *n* = 9 brain slices, 3 mice; *P*<0.0001, unpaired *t*-test). **(F and G)** Reduction of VGAT puncta per AIS in ChC-ablated mice (CTRL: *n* = 52 cells, 3 mice; DT: *n* = 57 cells, 3 mice; *P*<0.0001, unpaired *t*-test). **(H and I)** Reduction of gephyrin puncta per AIS in ChC-ablated mice (CTRL: *n* = 224 cells, 3 mice; DT: *n* = 243 cells, 3 mice; *P*<0.0001, unpaired *t*-test). Data are represented as mean ±SEM. \*\*\*\**P*<0.0001

To genetically ablate ChCs, we used intersectional Tau^loxP-STOP-loxP-FRT-STOP-FRT-DTR^ (*Tau-ds-DTR*) mice (*35*), in which the human diphtheria toxin receptor (DTR) gene is driven by the *Tau* promoter (Figure 1A). DTR expression is not activated until the removal of both of the STOP cassettes by Cre and Flp DNA recombinase. By combining the *Tau-ds-DTR line* with *Unc5b-CreER:Nkx2.1-Flp:Ai65* line, specific expression of DTR together with tdTomato in ChCs could be achieved. To determine the efficiency of genetic ablation of ChCs, *Unc5b-CreER:Nkx2.1-Flp:Tau-ds-DTR^+/-^:Ai65* mice were given TMX at P21 and P23 to turn on the expression of DTR and tdTomato in ChCs. Two weeks after TMX induction, mice were intraperitoneally (i.p.) injected with DT (50 μg/kg) at P37 and P39 to ablate the ChCs. The efficiency of genetic ablation was analyzed one week after DT application (Figure 1A). We found a marked reduction in ChC cell number and their innervation to the AIS (Figure 1B and 1C). There was an 81.06% reduction in tdTomato-positive cells in the PrL and IL regions of the dmPFC in the DTR-expressing mice compared with their littermate control (Figure 1B and 1D). Meanwhile, compared with 76.08% of PyN AISs were innervated by tdTomato-positive cartridges in Layer2/3 of dmPFC in the control littermates, only 21.77% of the AISs were innervated in the DTR-expressing mice, suggesting that the ablation of ChCs diminished their innervation on the pyramidal neuron AIS (Figure 1C and 1E). Furthermore, to assess whether the ablation of ChCs resulted in overall reduction of GABAergic synaptic input on the AIS, we analyzed the number of VGAT and gephyrin puncta at the AIS. We found that both VGAT and gephyrin puncta at the AIS were markedly reduced in the DTR-expressing mice compared to their littermates (Figure 1F to 1I). Therefore, we established an efficient way to reduce GABAergic synapses at the AIS of PyNs through the ablation of ChCs, which enabled us to explore whether and how axo-axonic GABAergic synaptic input contributes to AIS physiology.

### Reduction of AIS GABAergic synapses results in homeostatic changes in AIS morphology and physiology

To characterize the impact of ChC ablation on AIS properties, we analyzed the AIS of L2 PyNs in the dmPFC as they are densely innervated by ChCs (Figure S1). Using immunohistochemical staining for Ankyrin-G, the master organizer of the AIS molecular assembly and an granted AIS marker for morphological analysis (*36*), we observed that the length of the AIS was significantly shortened in dmPFC layer2 of the DTR-expressing mice compared to their littermates controls (Figure 2A and 2B), with no detectable changes regarding their width or the distance to cell body (Figure S3). Next, we examined the distribution of voltage-gated Na^+^ (Na_v_) channels as they are the most relevant channels clustered on the AIS that contribute to action potential initiation. Immunostaining of Pan Na_v_1 revealed that the expression level of the Na_v_1 family was significant reduced after the ablation of ChCs (Figure 2C-2E). Among the Na_v_1 family members, Na_v_1.2 and Na_v_1.6 are the two major members with enriched expression on the AIS of PyNs (*37, 38*). With the lowest activation threshold, Na_v_1.6 has been shown to contribute to the initiation of the AP (*5, 39*). Therefore, we further examined the expression of Na_v_1.6 on the AIS after ablating ChCs. We found that the expression of Na_v_1.6 was significantly reduced in the DT treated mice compared to their littermate controls (Figure 2F and 2G). However, the distribution of Na_v_1.6 along the AIS was not changed, remaining enriched at the distal AIS (Figure 2H).

**Fig. 2.**
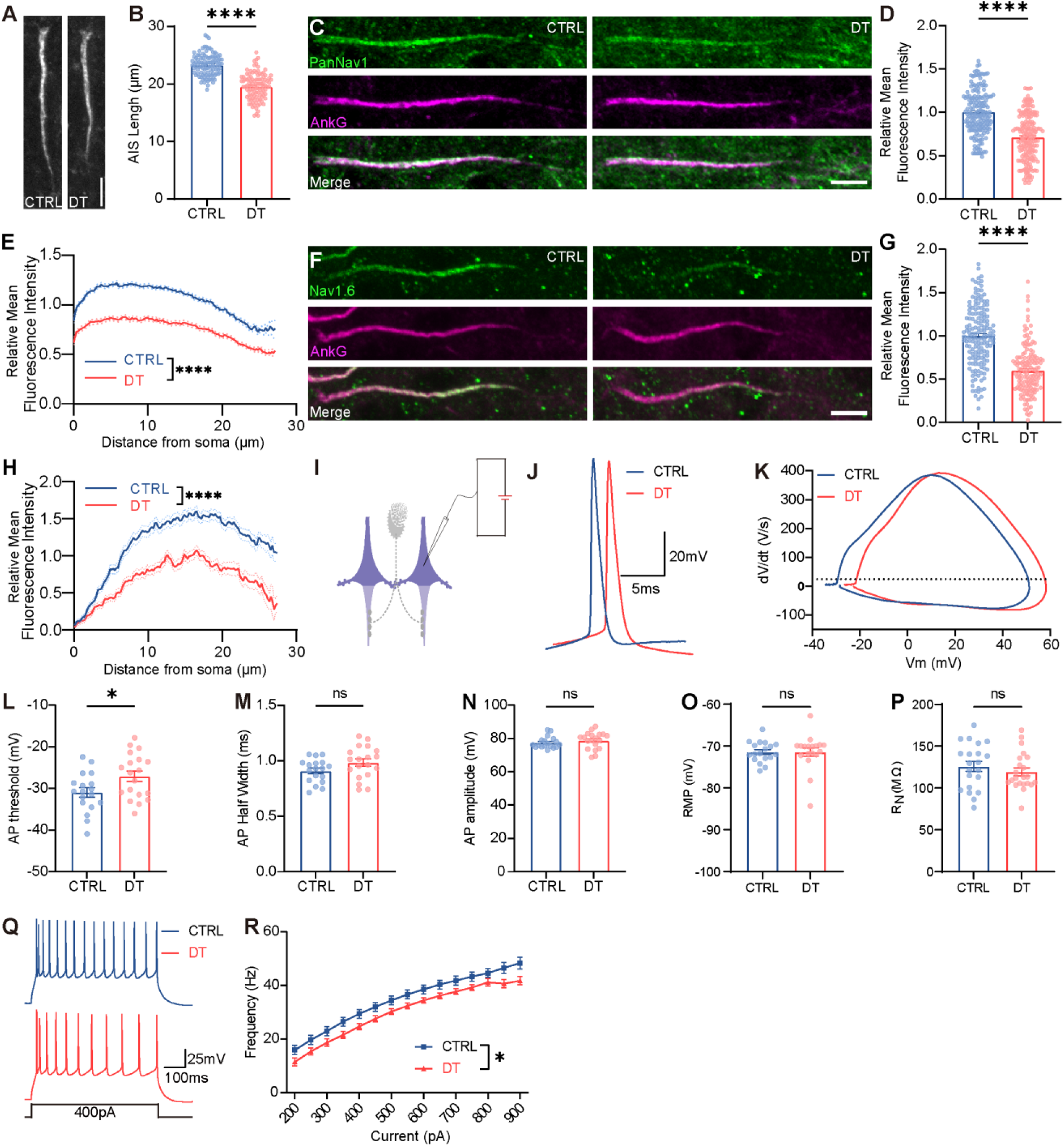
Ablation of ChCs induced AIS structural changes and decreases neuronal excitability. **(A)** Representative images of L2 PyN AIS without (CTRL) or with (DT) ChC ablation. Scale bar: 5 μm. **(B)** Reduced PyN AIS length in DT group compared to control group (CTRL: *n* = 224 cells, 3 mice; DT: *n* = 243 cells, 3 mice; *P*<0.0001, unpaired *t*-test). **(C)** Representative images of AIS with pan-Nav1 staining. **(D)** Decreased mean fluorescence intensity of pan-Nav1 after ChC ablation (CTRL: *n* = 173 cells, 3 mice; DT: *n* = 178 cells, 3 mice; *P*<0.0001, unpaired t-test). **(E)** Normalized mean fluorescence intensity of pan-Nav1 along AISs of both control and DT-treated group (CTRL: *n* = 173 cells, 3 mice; DT: *n* = 178 cells, 3 mice; *P*<0.0001, two-way ANOVA). **(F)** Representative images of AIS with Nav1.6 staining. **(G)** Decreased mean fluorescence intensity of Nav1.6 after ChC ablation (CTRL: *n* = 159 cells, 3 mice; DT: *n* = 147 cells, 3 mice; *P*<0.0001, unpaired *t*-test). **(H)** Normalized mean fluorescence intensity of Nav1.6 along AISs of both control and DT-treated group (CTRL: *n* = 159 cells, 3 mice; DT: *n* = 147 cells, 3 mice; *P*<0.0001, two-way ANOVA). **(I)** Schematic illustrating whole-cell patch recording of L2 PyNs that were innervated by ChCs. **(J)** Representative AP waveform of PyNs in control and DT-treated animals. **(K)** Phase plot of the AP waveform in **(J)**. **(L)** Increased AP threshold after ChC ablation by DT (CTRL: *n* = 18 cells, 5 mice; DT: *n* = 18 cells, 6 mice; *P* = 0.018, unpaired *t*-test). **(M to P)** No significant changes in AP half width **(M)**, amplitude **(N)**, RMP **(O)**, and R_N_ **(P)** between control and DT-treated group (CTRL: *n* = 18 cells, 5 mice; DT: *n* = 18 cells, 6 mice; *P* = 0.063 for half width, *P* = 0.411 for amplitude, *P* = 0.983 for RMP, *P* = 0.395 for R_N_, unpaired *t*-test). **(Q)** Representative traces of AP trains elicited by current injection. **(R)** Input/frequency relationship as determined by injections of increasing currents. The DT-treated group showed significantly decreased firing frequency (CTRL: *n* = 32, 5 mice; DT: *n* = 26, 6 mice. two-way ANOVA, *P* = 0.032). Data are represented as mean ±SEM. n.s. *P*>0.05, \**P*<0.05, \*\*\*\**P*<0.0001.

To examine whether the ablation of GABAergic synapses on the AIS was associated with electrophysiological changes, we performed whole-cell patch recordings in L2 PyNs of the dmPFC (Figure 2I). We observed an increased AP threshold, as illustrated by the AP phase plot (Figure 2J-2L), with no change in AP half-width and amplitude (Figure 2M and 2N), suggesting a decrease in neuronal excitability. Indeed, while there were no detectable changes in resting membrane potential (RMP) and input resistance (R_N_) (Figure 2O and 2P), the excitability of dmPFC L2 PyNs in DTR-expressing mice was significantly decreased (Figure 2Q and 2R).

Taken together, the ablation of ChCs and the resulting reduction in GABAergic synaptic input to the AIS induced the shortening of the PyN AIS, a reduction in Na_v_1 expression, and an increase in AP threshold. These plastic changes in AIS properties are in line with the observed decrease in neuronal excitability.

### Blocking axo-axonic synaptic transmission induces similar changes in AIS as ablating ChC

Although genetic ablation of ChCs provided a valuable tool to study the function of axo-axonic synapse, global cell loss might trigger microglia activation and neuroinflammation, introducing undesirable artifacts (*40*). To validate our findings, we locally injected an adeno-associated virus (AAV) encoding a Cre recombinase-dependent tetanus toxin (DIO-Tettox) into the dmPFC of *Unc5b-CreER* mice to specifically block synaptic transmission of L2 ChCs (Figure 3A). As AAV9 has limited tropism toward capillary endothelial cells (*41*), we did not identify any endothelial cells infected by the AAV we used in this study. To avoid any batch effects, we stereotaxically injected DIO-Tettox-2A-mCherry virus into the dmPFC of one hemisphere and injected DIO-tdTomato control virus contralaterally in the same animal (Figure 3B). Then, we introduced i.p. injection of TMX two days after viral injection to induce recombination. To track the time course when chronic inhibition from blocking axo-axonic synaptic transmission exerts its impact on AIS physiology, we performed immunostaining against Ankyrin-G (AnkG) and Na_v_1.6 at three time points: 2 weeks, 3 weeks and 4 weeks post-viral injection (Figure 3C). Interestingly, we found that the length of the AIS started to change after 3 weeks post-injection (Figure 3D), and the expression level of Na_v_1.6 did not change until 4 weeks post-injection (Figure 3E). These data indicated that reduced ChC synaptic transmission was sufficient to induce homeostatic changes in AIS properties.

**Fig. 3.**
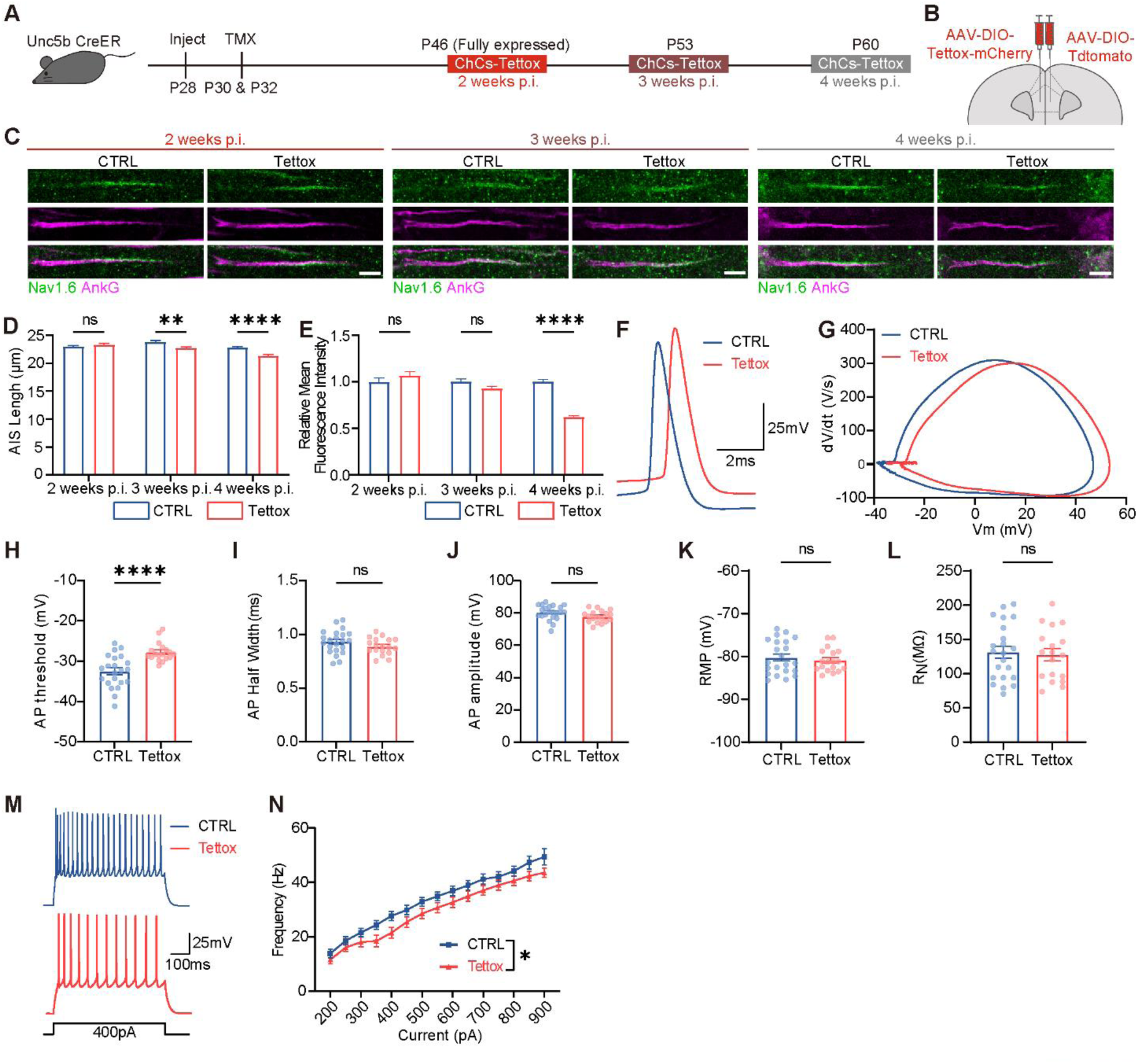
Inactivation of axon-axonic neurotransmission leads to homeostatic changes of the PyN AISs in dmPFC. (**A**) Schematic showing the strategy of blocking axon-axonic transmission in dmPFC. TMX was applied at P30 and P32 to induce tetanus toxin (Tettox) expression in ChCs. Immunohistochemistry was performed at 2 weeks, 3 weeks, and 4 weeks post-injection (p.i.). **(B)** Schematic illustrating the unilateral injection of AAV-DIO-Tettox-mCherry and AAV-DIO-tdTomato in the same animal. **(C)** Representative images of immunostaining against Nav1.6 and AnkG at the indicated phases. Scale bar: 5 μm. **(D)** Relative mean fluorescence intensity of Na_v_1.6 in PyN AISs in the Tettox group compared to the control group at the indicated phases. **(E)** PyN AIS length in the Tettox group compared to the control group at the indicated phases (2 weeks p.i.: *n* = 101 cells from 3 mice for CTRL group, *n* = 62 cells from 3 mice for Tettox group, *P* = 0.278 for AIS length, *P* = 0.276 for fluorescence intensity; 3 weeks p.i.: *n* = 92 cells from 3 mice for CTRL group, *n* = 114 cells from 3 mice for Tettox group, *P* = 0.007 for AIS length, *P* = 0.105 for fluorescence intensity; 4 weeks p.i.: *n* = 121 cells from 3 mice for CTRL group, *n* = 121 cells from 3 mice for Tettox group, *P* < 0.0001 for AIS length, *P* < 0.0001 for fluorescence intensity. Unpaired Holm-Sidak multiple *t*-test). **(F)** Representative AP waveform of PyNs in control and Tettox-treated animals. **(G)** Phase plot of the AP waveform in **(F)**. **(H)** Increased AP threshold after blocking axon-axonic synaptic transmission by Tettox (CTRL: *n* = 22 cells, 4 mice; Tettox: *n* = 18 cells, 4 mice; *P* < 0.0001, unpaired *t*-test). **(I to L)** No significant changes in AP half width **(I)**, amplitude **(J)**, RMP **(K)**, and R_N_ **(L)** between the control and Tettox group (CTRL: *n* = 22 cells, 4 mice; Tettox: *n* = 18 cells, 4 mice; *P* = 0.168 for half width, *P* = 0.066 for amplitude, *P* = 0.528 for RMP, *P* = 0.771 for R_N_, unpaired *t*-test). **(M)** Representative traces of AP trains elicited by current injection. **(N)** Input/frequency relationship as determined by injections of increasing currents. The Tettox group showed significantly decreased firing frequency (CTRL: *n* = 26, 4 mice; Tettox: *n* = 26, 4 mice, *P* = 0.042, two-way ANOVA). Data are represented as mean ±SEM. n.s. *P*>0.05, \**P*<0.05, \*\*\*\**P*<0.0001

To corroborate that the structural and ion channel changes of the AIS are coupled with the functional changes, we performed whole-cell patch recording in L2 PyNs in the viral injection sites 4 weeks post-injection. Consistent with the effects of ChC ablation, the AP threshold was significantly increased in the Tettox group compared with the control group (Figure 3F-3H), with no change in AP half-width or amplitude (Figure 3I and 3J). With no change in RMP and R_N_ (Figure 3K and 3L), the AP firing rate induced by the injection of stepped currents was also reduced, suggesting that the excitability of PyNs was decreased when chronically blocking axo-axonic synaptic transmission (Figure 3M and 3N).

In summary, chronic reduction of axo-axonic input leads to shortened AIS, decreased Na_v_1.6 expression, increased AP threshold, and decreased intrinsic excitability of PyNs, which is consistent with the phenotypes caused by genetic ablation of ChCs.

### Chronic activation of ChCs elicits homeostatic plasticity of the PyN AIS

We next explore whether chronic activation of ChCs might play a role in the plasticity of the AIS. We used designer receptors exclusively activated by designer drugs (DREADDs), hM3Dq, to increase the axo-axonic synaptic activity (*42*). We virally expressed Cre-dependent DIO-hM3Dq or DIO-EGFP in each of the hemispheres of the *Unc5b-CreER* mouse, respectively, followed by TMX induction 2 days later (Figure 4A and 4B). After 2 weeks, CNO was applied twice a day for one week to elicit chronic activation of ChC (Figure 4A). Interestingly, chronic activation of ChCs resulted in a significant elongation of the PyN AISs and an increase in Na_v_ 1.6 expression (Figure 4C-4E). In parallel to this, by whole-cell patch recoding (Figure 4F), we found that the PyN exhibited a decrease in AP threshold (Figure 4G-4I) with unchanged AP half-width and amplitude (Figure 4J and 4K). With no detectable changes in their RMP and R_N_ (Figure 4L and 4M), the neuronal excitability of the L2 PyNs in the dmPFC was significantly increased (Figure 4N and 4O).

**Fig. 4.**
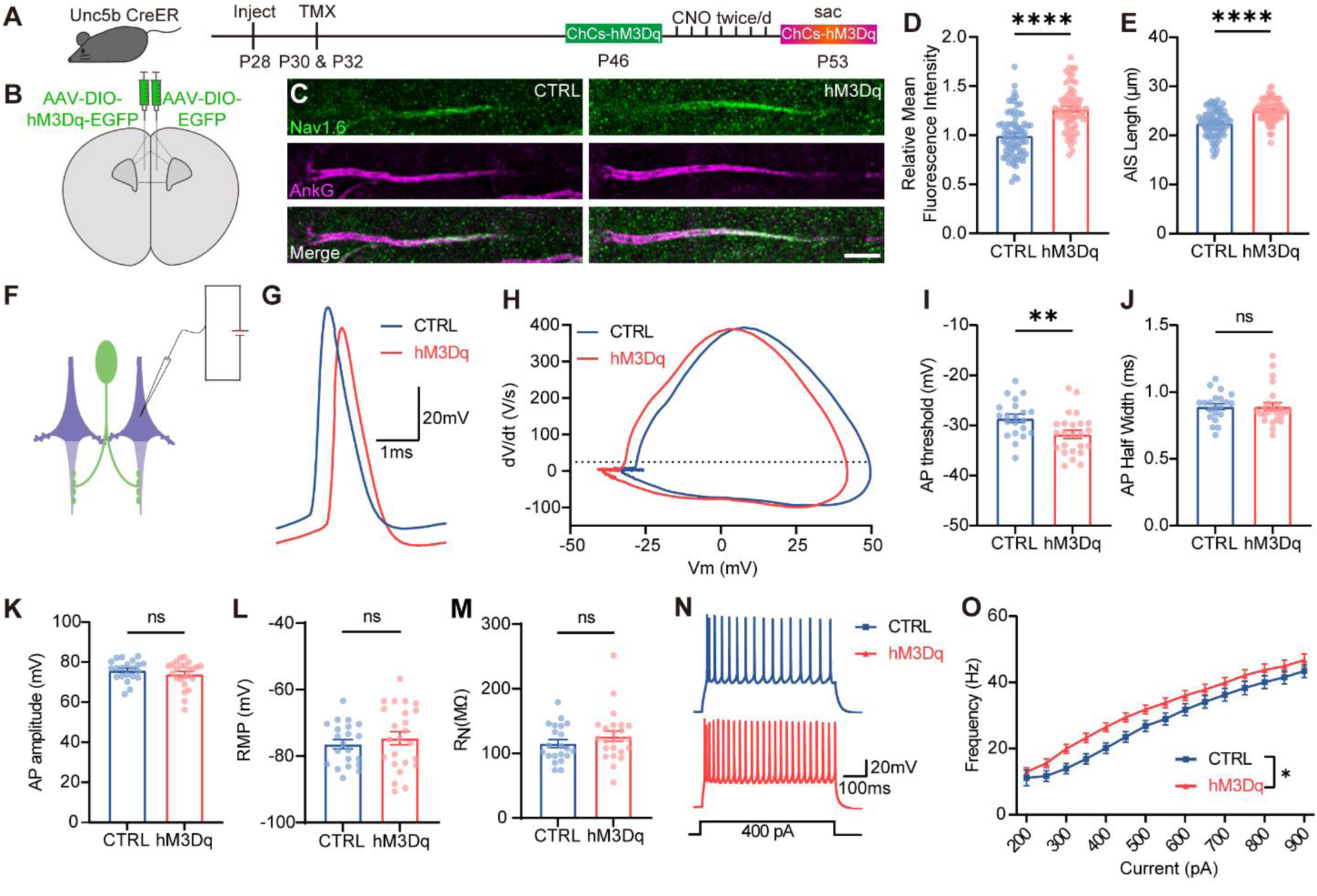
Chronic pharmacogenetic activation of ChCs evoked homeostatic plasticity of PyN AIS. (**A**) Schematic showing the strategy of chronic activation of ChCs. TMX was applied at P30 and P32 to induce hM3Dq expression in ChCs. CNO (0.6mg/kg) was applied twice daily during P46-53. Animals were sacrificed to investigate the structure of the PyN AIS at P53. **(B)** Schematic illustrating the unilateral injection of AAV-DIO-hM3Dq-EGFP and AAV-DIO-EGFP in each hemisphere. **(C)** Representative images of immunostaining against Nav1.6 and AnkG. Scale bar: 5 μm. **(D)** Increased mean fluorescence intensity of Nav1.6 after chronic activation of ChCs (CTRL: *n* = 83 cells, 3 mice; hM3Dq: *n* = 68 cells, 3 mice; *P* < 0.0001, unpaired *t*-test). **(E)** Increased PyN AIS length in the hM3Dq group compared to the control group (CTRL: *n* =83 cells, 3 mice; hM3Dq: *n* = 68 cells, 3 mice; *P* < 0.0001, unpaired *t*-test). **(F)** Schematic illustrating whole-cell patch recording of L2 PyNs innervated by hM3Dq-expressed ChCs. **(G)** Representative AP waveform of PyNs in control and pharmacogenetically activated ChCs animals. **(H)** Phase plot of the AP waveform in **(G)**. **(I)** Decreased AP threshold after chronic activation of ChCs (CTRL: *n* = 21 cells, 5 mice; hM3Dq: *n* = 24 cells, 5 mice; *P* = 0.009, unpaired *t*-test). **(J to M)** No significant changes in AP half-width **(J)**, amplitude **(K)**, RMP **(L)**, and R_N_ **(M)** between the control and hM3Dq group (CTRL: *n* = 21 cells, 5 mice; hM3Dq: *n* = 24 cells, 5 mice; *P* = 0.996 for half width, *P* = 0.290 for amplitude, *P* = 0.472 for RMP, *P* = 0.276 for R_N_, unpaired *t*-test). **(N)** Representative traces of AP trains elicited by current injection. **(O)** Input/frequency relationship as determined by injections of increasing currents. The hM3Dq group showed significantly increased firing frequency (CTRL: *n* = 28, 5 mice; Tettox: *n* = 34, 5 mice; *P* = 0.022, two-way ANOVA). Data are represented as mean ±SEM. n.s. *P*>0.05, \**P*<0.05, \*\**P*<0.01\*\*\*\**P*<0.0001

Taken together, we found that increased activation of ChCs for one week caused elongation of the AIS and increased expression of Na_v_1.6 at the AIS. These morphological and molecular changes were coupled with a decreased AP threshold and increased intrinsic excitability of PyNs. Therefore, chronic alterations of ChC activity bidirectionally induced homeostatic changes in PyN excitability by tuning the morphology and Na_v_1.6 expression at the AIS.

### Enhancement of somatic inhibition does not elicit structural changes of the AIS

We next sought to investigate whether other forms of inhibitory input might induce the same homeostatic changes in the AIS. Parvalbumin (PV) fast-spiking interneurons are the main source of perisomatic inhibition in the cortex (*43, 44*). Traditionally, ChCs and PV basket cells are thought to be the two major subtypes of PV fast-spiking interneurons, with ChCs targeting the AIS and PV basket cells targeting the soma (*45*). Therefore, we set out to investigate whether chronic activation of PV basket cells might have any effect on AIS plasticity. Although both ChCs and basket cells are believed to express PV as a biomarker, recent single-cell RNA sequencing data have revealed that ChCs can be further grouped into PV-high and PV-low subtype (*32*). Therefore, we set out to determine the properties of ChCs in the dmPFC. First, we crossed the *PV-Flp* driver line with the *Unc5b-CreER* driver line and the intersectional reporter line, *Ai65,* to label the PV-positive ChCs (*Unc5b-CreER:PV-Flp:Ai65*). Interestingly, few tdTomato-positive cells were shown in the dmPFC, suggesting that ChCs in the dmPFC are the PV-low subtype (Figure 5A). To corroborate this observation, we performed immunohistochemistry with PV antibody in the *Unc5b-CreER:Nkx2.1-Flp:Ai65* brain slices. Consistent with what we found in the *Unc5b-CreER:PV-Flp:Ai65* line, the expression level of PV was hardly detectable in the ChCs in the dmPFC region, confirming that ChCs in the dmPFC are the PV-low subtype (Figure 5B). These findings provide us with an opportunity to differentially manipulate PV basket cells and ChCs in this area, thus further dissecting their functional differences.

**Fig. 5.**
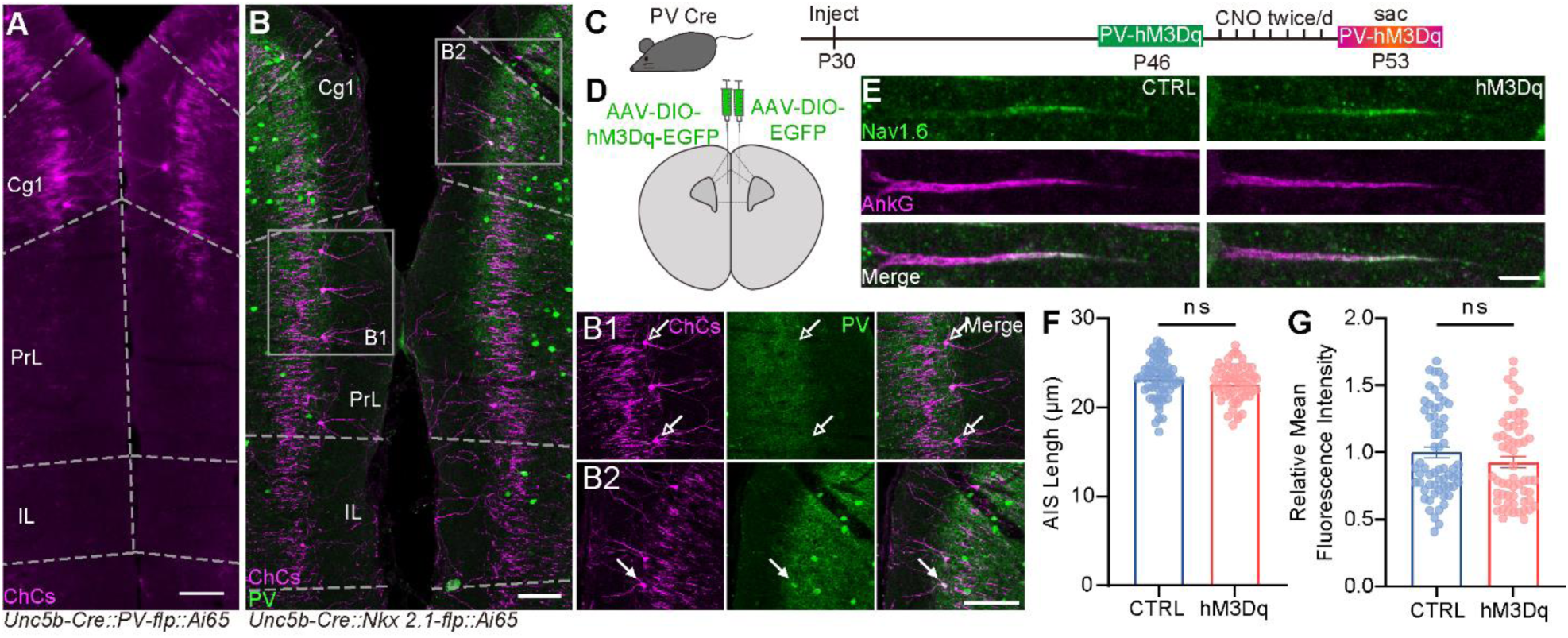
Chronic activation of somatic inhibition failed to induce the plastic changes of the AIS. (**A**) Representative image showing that few ChCs were labeled in dmPFC of *Unc5b-CreER; PV-Flp; Ai65* mouse. Scale bar: 500 μm. **(B)** Immunostaining of PV in the *Unc5b-CreER; Nkx2.1-Flp; Ai65* mouse. Scale bar: 100 μm. Insets: hollow arrows indicate ChCs with little PV immunosignals; solid arrows indicate ChCs co-localize with PV immunosignals. Scale bar: 100 μm. **(C)** Schematic showing the strategy of chronic activation of somatic inhibition. CNO (0.6mg/kg) was applied twice daily from P46 to P53. Animals were sacrificed at P53. **(D)** Schematic illustrating the injection of AAV-DIO-hM3Dq-EGFP and AAV-DIO-EGFP in the same animal. **(E)** Representative images of AIS stained with Na_v_1.6 and AnkG antibodies. Scale bar: 5 μm. **(F)** No significant change in the mean fluorescence intensity of Na_v_1.6 after chronic activation of somatic inhibition (CTRL: *n* = 63 cells, 3 mice; hM3Dq: *n* = 58 cells, 3 mice; *P* = 0.174, unpaired *t*-test). **(G)** No significant change of the AIS length in the control group and hM3Dq group (CTRL: *n* = 63 cells, 3 mice; hM3Dq: *n* = 58 cells, 3 mice; *P* = 0.228, unpaired *t*-test). Data are represented as mean ±SEM. n.s. *P*>0.05.

To determine whether chronic activation of PV basket cells had any effect on AIS plasticity, we virally expressed Cre-dependent DIO-hM3Dq or DIO-EGFP in each hemisphere of the dmPFC region in the same *PV-Cre* mouse (Figure 5C and 5D). Two weeks later, CNO (0.6mg/kg) was applied intraperitoneally for 7 days to achieve chronic activation of PV basket cells (Figure 5C). As shown in Fig 5E, chronic activation of PV basket cells for 7 days did not alter the length of the AIS of PyNs in the dmPFC region (Figure 5F), nor did it change the expression level of Na_v_1.6 at the AIS (Figure 5G). These results indicate that chronic alteration of somatic inhibition from PV basket cells is not sufficient to induce plastic changes in the PyN AIS, signifying a requirement of direct local synaptic input from ChCs.

### Behavioral deficits caused by blocking ChC transmission is reversed when AIS plasticity occurs

To further explore the physiological and functional implications of AIS homeostatic plasticity in the context of altered axo-axonic synaptic transmission, we took advantage of the time course of homeostatic changes by virally blocking axo-axonic synaptic transmission in the dmPFC (Figure 3A-D). We first assessed behavioral deficits 2 weeks after injection of AAV-DIO-Tettox. Then, we performed the same behavioral test 4 weeks after injection to assess any changes that might be associated with AIS plasticity (Figure 6A). We started with the open field test to examine exploratory behavior and locomotor activity, and found no deficits at either time points post-injection (Figure S4). Similarly, the mice did not show any anxiety-related behavior in the elevated plus maze test (Figure S4). Given the large number of studies showing that the dmPFC network is critical for higher-order brain functions such as social behavior (*46, 47*), we further applied the three-chamber social test to evaluate social ability and social novelty (Figure 6B). At 2 weeks post-injection, we found that the Tettox-expressing group showed less interest in socializing with stranger mice (Figure 6C-6F). The ratio of interaction time in the stranger box versus object box was reduced in the Tettox group compared with control group (Figure 6D and 6E). Furthermore, the total social time was significantly less in the Tettox group (Figure 6F). However, in the social novelty test, the Tettox-expressing mice were equally driven to the novel stranger as the control mice (Figure 6D). These data indicated that inhibition of ChC transmission in the mPFC led to a deficit in social ability. Next, we evaluated the effects of chronic inhibition of ChC transmission on social behavior. Unexpectedly, the Tettox group showed increased sociability compared with the control group at 4 weeks post-injection (Figure 6G-6J). The ratio of interaction time in the stranger box versus object box was also increased in the Tettox group (Figure 6H and 6I), whereas the total social time did not differ between the two groups (Figure 6J). These data indicate that the deficits in sociability observed at 2 weeks post-injection were reversed at 4 weeks post-injection.

**Fig. 6.**
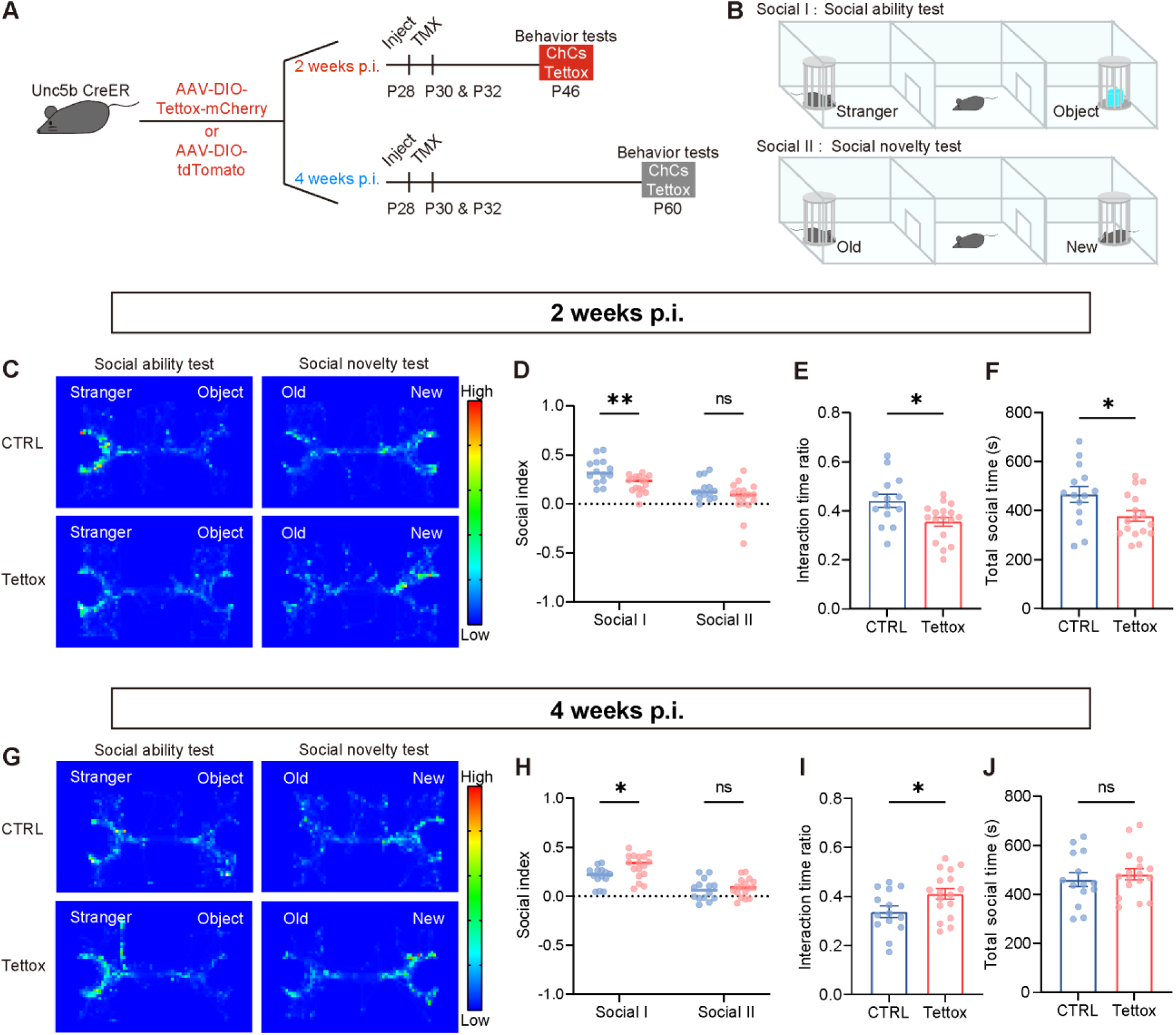
Effect of reduced axon-axonic neurotransmission on social behavior. (**A**) Schematic showing the strategy of acute and chronic blocking of the dmPFC ChC axon-axonic transmission. TMX was applied at P30 and P32 to induce Tettox expression in ChCs. Two weeks after TMX induction (P46) was defined as acute inhibition and 4 weeks after TMX induction (P60) was defined as chronic inhibition. **(B)** Schematic illustrating the apparatus of 3-chamber social test. **(C)** Representative heatmaps of indicated groups in the acute inhibition condition. **(D)** Reduced preference in social approach but not social novelty test in the Tettox group in the acute condition (*n* = 14 for CTRL, *n* = 17 for Tettox, *P* = 0.005 for social ability, *P* = 0.121 for social novelty, unpaired Holm-Sidak multiple *t-*test). **(E-F)** Reduced interaction time ratio **(E)** and total social time **(F)** in Tettox group in the acute condition (*n* = 14 for CTRL, *n* = 17 for Tettox, *P* = 0.012 for exploration time ratio, *P* = 0.028 for total social time, unpaired *t*-test). **(G)** Representative heatmaps of indicated groups in the chronic inhibition condition. **(H)** Increased preference in social approach but not social novelty test in the Tettox group in the chronic condition. (*n* = 14 for CTRL, *n* = 17 for Tettox, *P* = 0.034 for social ability, *P* = 0.575 for social novelty, unpaired Holm-Sidak multiple *t-*test) **(I)** Increased interaction time ratio in the chronic Tettox group. (*n* = 14 for CTRL, *n* = 17 for Tettox, *P* = 0.033, unpaired *t*-test) **(J)** No change in total social time in the Tettox Group in the chronic condition (*n* = 14 for CTRL, *n* = 17 for Tettox, *P* = 0.569, unpaired *t*-test). Data are represented as mean ±SEM. n.s. *P*>0.05, \**P*<0.05, \*\**P*<0.01.

We next explored whether the AIS plasticity tuned by enhancement of ChC activity was correlated with changes in social behavior as well. We bilaterally injected AAV-DIO-hM3Dq or DIO-EGFP (as control) into the dmPFC of *Unc5b-CreER* mice, followed by TMX induction 2 days later. The hM3Dq was allowed to be expressed for 2 weeks. First, we tested whether acute activation of dmPFC ChCs would elicit any changes in social behavior. CNO was applied to animals 30 minutes before the behavioral test to achieve acute activation (Figure 7A). We found that mice with acute enhancement of ChC activity displayed a significant increase in the time spent in the social box, indicating that activation of ChCs had a prosocial effect (Figure 7B-7E). The saline (SAL) groups were also included as control groups, and no significant difference was found between the EGFP-SAL group and hM3Dq-SAL group (Figure S5). Next, we examined whether the timing of AIS plasticity was coupled with changes in social behavior. We performed the social test after chronic activation of ChCs. Specifically, CNO was injected intraperitoneally twice daily to induce chronic enhancement of axon-axonic synaptic transmission. After 7 consensus days, when the AIS had elongated, we performed social tests (Figure 7A). Unlike acute activation, mice with chronic activation of ChCs showed normal social ability compared with the control group (Figure 7F-7I). The interaction time ratio of stranger mice and the total social time showed no difference between the chronic activation group and the control group (Figure 7H and 7I). Similarly, the social novelty test showed no significant difference between the chronic activation group and the control group (Figure 7G).

**Fig. 7.**
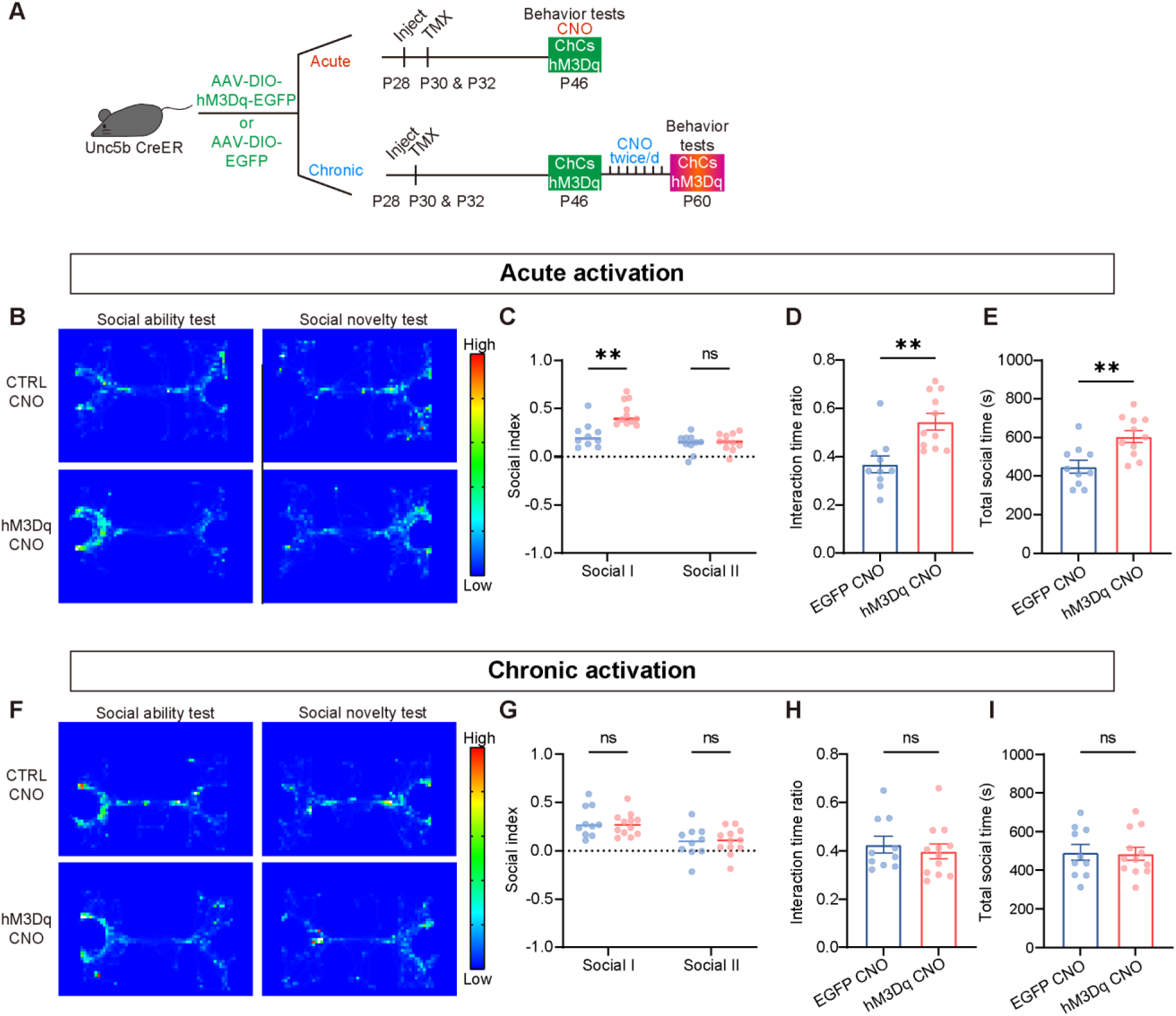
Effect of pharmacogenetic activation of dmPFC ChCs on social behavior. (**A**) Schematic showing the strategy of acute and chronic pharmacogenetic activation of the dmPFC ChC axon-axonic transmission. AAV virus encoding DIO-hM3Dq-EGFP or DIO-EGFP was injected bilaterally at P28, TMX was applied at P30 and P32. Two weeks after TMX induction (P46) was defined as acute inhibition and 4 weeks after TMX induction (P60) was defined as chronic inhibition. **(B)** Representative heatmaps of animals expressing AAV-DIO-EGFP or AAV-DIO-hM3Dq with acute activation of ChCs. **(C)** Increased preference in social approach but not social novelty test in the hM3Dq group after acute activation of ChCs (*n* = 10 for EGFP CNO, *n* = 11 for hM3Dq CNO, *P* = 0.001 for social ability, *P* = 0.704 for social novelty, unpaired Holm-Sidak multiple *t*-test). **(D-E)** Increased interaction time ratio **(D)** and total social time **(E)** in hM3Dq group after acute activation of ChCs (*n* = 10 for EGFP CNO, *n* = 11 for hM3Dq CNO. In **(D)**, *P* = 0.002, in **(E)**, *P* = 0.003, unpaired *t*-test) **(F-I)** No difference between CTRL and hM3Dq group in social approach or social novelty test in the hM3Dq group after chronic activation of ChCs (*n* = 10 for EGFP CNO and 12 for hM3Dq CNO. In **(G)**, *P* = 0.885 for social ability, *P* = 0.904 for social novelty, unpaired Holm-Sidak multiple *t*-test. In **(H)**, *P* = 0.571, unpaired *t*-test. In **(I)**, *P* = 0.904, unpaired *t*-test., unpaired *t*-test). Data are represented as mean ±SEM. n.s. *P*>0.05, \*\**P*<0.01.

Taken together, our findings suggest that acute and chronic alterations of axon-axonic synaptic input lead to different outcomes in social behavior (Figure 8). The fact that the deficits in social ability recovered when homeostatic plasticity of the AIS occurred insinuates a link between AIS plasticity and behavioral recovery. However, the excessive sociability of the Tettox group suggested an over-compensatory effect, indicating the importance of an intact microcircuit in maintaining normal behavior.

**Fig. 8.**
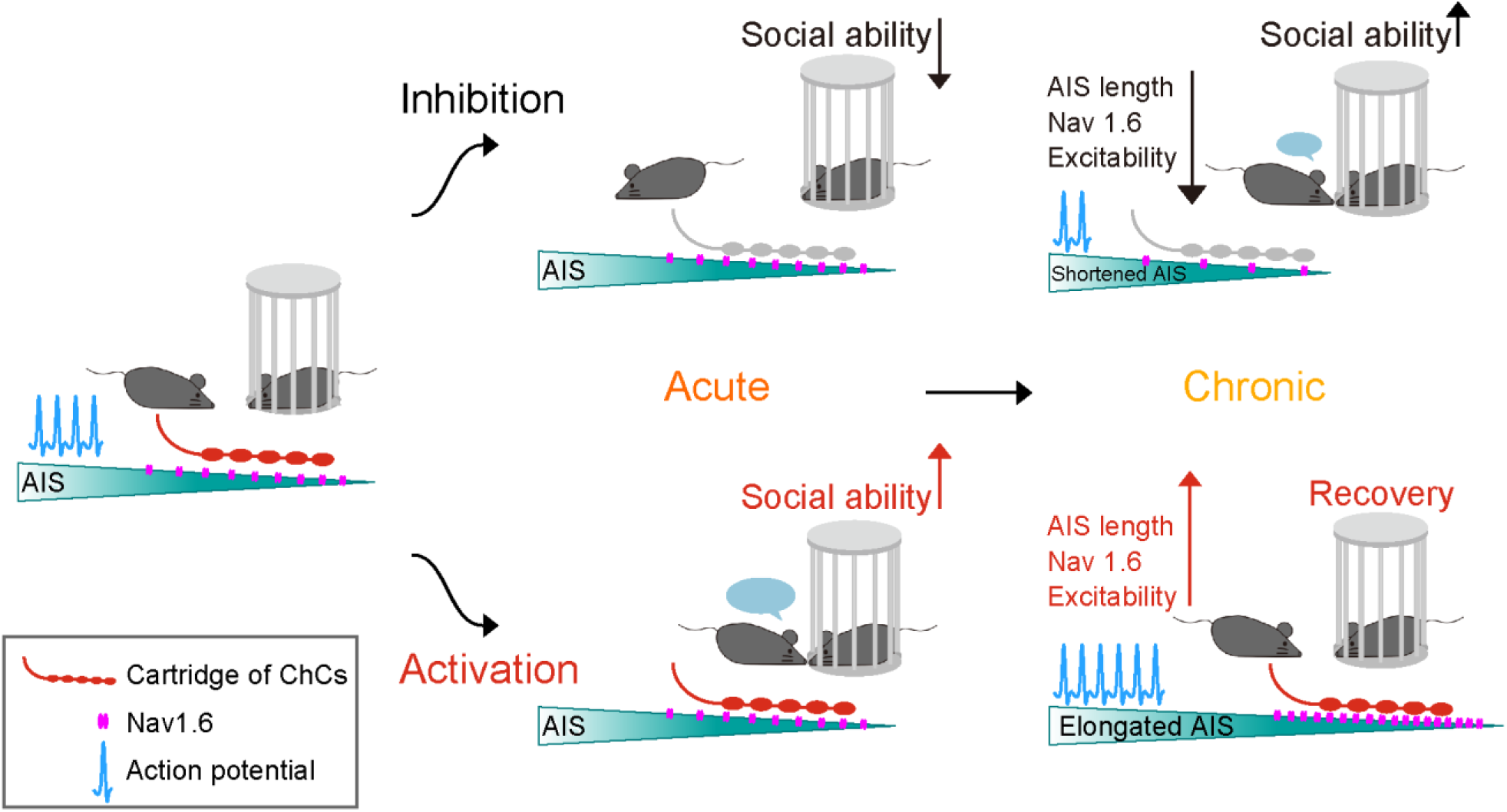
Schematics illustrating sociability changes induced by alterations of ChC synaptic input are recovered when homeostatic plasticity of the AIS occurs.

## Discussion

Previous studies have shown that alterations of network activity cause homeostatic rearrangement of PyN AIS(*11, 12, 19*). Plastic changes of the AIS effectively adjust the properties of AP firing and propagation, which in turn oppose ongoing deviations of activity levels and stabilize network functions. Here, we extended these studies by investigating AIS plasticity responding to changes in ChC synaptic input. We found that chronic alterations of synaptic input to the AIS by blocking axo-axonic synaptic transmission or by pharmacological enhancement of ChCs induced structural and functional changes of the AIS. Altered AIS structure and expression levels of voltage-gated sodium channel correlated with shifted initiation thresholds of AP, which were reflected by changes in neuronal excitability. The time course of homeostatic tuning of AIS structure and neuronal excitability in dmPFC caused by altering ChC synaptic transmission correlates with changes in social behavior. Our results provide evidence for input-specific regulation of AIS plasticity that contributes to compensatory changes in disturbed functional microcircuits.

Recent work has demonstrated that the synaptic plasticity of ChCs follows a homeostatic rule when network activity changes(*48*). Modulation of PyN activity in the somatosensory cortex effectively adjust the innervation of ChCs to the PyN AISs to stabilize network activity. Interestingly, axo-axonic synapse plasticity can be driven in a cell-autonomous manner. Pharmacological activation of ChCs in early developmental stages (P12-P18, when GABA is depolarizing) induces a decrease in synaptic number and connection probability. Although there is no significant change in AIS length of the innervated PyNs, a trend of reduction is observed(*48*). Typically, one PyN AIS receives 1-4 cartridges from different ChCs(*49*). Therefore, sparsely infected single ChCs, by utilizing *Nkx2.1-CreER* line, might be insufficient to drive the homeostatic changes of the PyN AIS(*33*). In our study, we infect ChCs at higher density by using the *Unc5b-CreER* line(*25*). Manipulating axo-axonic synaptic activity with higher efficiency provided sufficient driving force to induce homeostatic plasticity of the AIS.

AACs are an interneuron subtype identified only in mammals so far and are located solely in the neocortex or allocortex, but not in other brain areas (*50*). Our findings that the AISs of cortical PyNs adapt to direct synaptic input to the AIS suggest that PyNs in the cortex might possess a strategic adjustment capacity that subcortical neurons lack. Our previous findings showed that dmPFC ChCs preferentially innervate PyNs projecting to basal lateral amygdala (_BLA_PyNs) over those projecting to the contralateral prelimbic cortex (_cPL_PyNs)(*27, 51*). In the territory of a single ChC axonal collateral, around 80% _BLA_PyNs are innervated, whereas around 60% for _cPL_PyNs are innervated (*51*). Meanwhile, cartridges on the AIS of _BLA_PyNs harbor significantly more boutons than those of on the _cPL_PyNs, indicating that the AISs of _BLA_PyNs receive more ChC input than those of _cPL_PyNs (*51*). This finding poses the possibility that the AISs of different cortical PyN subtypes might have varying degrees of response to local synaptic input. It is worth our future endeavor to further dissect the cell-type-specific features of AIS plasticity.

Interneurons can actively control the synaptic plasticity of cortical PyNs(*52–54*). Plasticity of local inhibitory circuits is critical to excitatory neuron plasticity during experience-dependent plasticity and learning(*55–59*). Subtype-specific alterations is believed to be able to control synaptic plasticity at subcellular resolution (*55*). However, prolonged changes in interneuron activity could trigger postsynaptic excitatory neuron hyperactive, and homeostatic regulation is necessary to safeguard proper circuit functions. Here we show that enhanced activation of ChCs, but not PV basket cells, induces the structural alteration of the AIS of excitatory neuron. It is important to note that, changes in global activity at a greater magnitude than activating locally infected PV basket cells, as those induced by adding high concentration of K^+^ in the *in vitro* neuronal cultures or altered sensory input (*11, 12, 19, 20*), could also induce plastic changes in the AIS. However, the fact that we did not observe a plastic change in AISs of PyN after manipulating PV basket cells in our experimental set up suggests that AIS-specific inputs may be a much more potent driver to induce structural and functional homeostatic changes in this subcellular domain. Such subcellular-domain specific homeostatic mechanism allows the cells to adjust themselves in a more sophisticated manner on top of those responding to more generic changes in network activity. Combined with the fact that ChCs and PV basket cells show significant differences in local and long-range inputs(*27, 60*), we speculate that homeostatic regulations operate not only under generic changes of network activity but also under activity changes in specific microcircuits. Previous reports have focused on how interneurons adapt network activity to act as local homeostats (*61, 62*), whereas our work suggests that interneuron subtype initiates plastic changes of PyN AIS, providing an appealing idea that homeostatic plasticity happens at the subcellular level when microcircuit activity is disturbed.

Intact dmPFC cortical neuronal excitatory and inhibitory balance is critical for higher cognitive functions(*63*). Compromises in dmPFC cortical inhibition are often found in animal models of autism or schizophrenia, which manifest social dysfunction (*46, 64, 65*). Although functional differentiation has been implicated among cortical interneuron subtypes in social behavior (*47*), acutely restoring E:I balance by either decreasing the excitability of excitatory neurons or increasing the excitability of PV neurons is sufficient to rescue social deficits in animal models of autism (*66*). While inhibition of somatostatin (SST) does not impair sociability, synchronization of either PV neurons or SST neurons improves sociability(*47*). Our observation that social impairment caused by blocking ChC synaptic transmission is restored when AIS tuning has occurred is in line with these reports that restoration of balanced network activity is sufficient to improve sociability. However, our results show increased sociability compared with control animals 4 weeks after injecting the Tet-tox AAV to ChC, suggesting overcompensation occurs to correct the loss of ChC input. Future studies are required to reveal the mechanistic relationships between compensatory activity of PyNs and the behavioral consequences.

The nervous system’s ability to adapt to various perturbations is crucial for maintaining normal brain functions. Our study has shown that social behavior is impaired after acutely blocking ChC synaptic transmission in the dmPFC. However, these behavioral deficits are reversed when AIS plasticity occurs, suggesting that intrinsic homeostatic plasticity is a key component for maintaining normal animal behavior. In line with our observation, several genetic mouse models of ASD have demonstrated impaired intrinsic homeostatic plasticity (*67–69*). In addition, studies have reported a decrease in ChC count, but not PV-basket cell count, in the mPFC region of individuals with ASD, further highlighting the critical role of ChC and AIS-homeostasis in its pathology (*70*). Nonetheless, further research is needed to determine whether the loss of AIS plasticity in response to altered ChC activity contributes to the etiology of ASD and other brain disorders.

Together, the results presented here support the idea that disturbance of subtype-specific inhibitory input can drive homeostatic plasticity of PyNs at the subcellular level. Prolonged alteration of axo-axonic synaptic input induced structural and functional plasticity of the AIS, which contributed to the tuning of neuronal excitability. Future studies are required to explore whether other subcellular structures of PyNs undergo homeostatic plasticity after the disturbance of subtype-specific inhibitory input and whether they contribute to stabilizing normal brain functions.

## Materials and Methods

### Animals

All procedures were conducted in accordance with the protocols approved by the Institutional Animal Care and Use Committee of Fudan University. *Unc5b-CreER* and *Nkx2.1-Flp* were provided by Z.J. Huang (Duke University School of Medicine), while *PV-Cre* (JAX Stock 008069), *PV-FlpE* (JAX Stock 021191), and *Rosa26-loxpSTOPloxp-frtSTOPfrt-TdTomato* (Ai65, JAX Stock 021875) were from Jackson Laboratory. *Tau-loxpSTOPloxp-frtSTOPfrt-DTR* (*Tau-ds-DTR*) was acquired from M. Goulding (Salk Institute). To specifically label ChCs, we combined *Unc5b-CreER* driver with *Nkx2.1-Flp* and the *Ai65* intersectional reporter. Ablation of ChCs was achieved by crossing *Unc5b-CreER:Nkx2.1-Flp:Ai65* with *Tau-ds-DTR.* Genetic labeling of PV-positive ChCs was done by crossing *Unc5b-CreER* and *PV-Flp* with *Ai65* reporter line. Wild-type C57BL/6j mice (from Charles River, Shanghai, China) were used as stranger mice in 3-chamber social test. All mice were maintained under standard conditions of 20 °C–24 °C, 40%–60% relative humidity, and a 12-hour light/dark cycle, with food and water freely available. The age of animals used ranged from P30 to P63. Mice of both sexes were used in all experiments except for the behavioral tests, in which only male mice were used.

### Tamoxifen induction

Tamoxifen (Sigma-Aldrich, MO, USA) was dissolved in corn oil (20 mg/ml) at room temperature by ultrasonication. Stocks were stored at 4°C for no more than 1 month. Tamoxifen was administered intraperitoneally twice, at a dose of 2 mg per 20 g body weight. For labeling or ablating ChCs, *Unc5b-CreER:Nkx2.1-Flp:Ai65* or *Unc5b-CreER:Nkx2.1-Flp:Ai65:Tau-ds-DTR* mice were injected at P21 and P23; for *Unc5b-CreER* mice infected with either AAV-tetanus toxin or AAV-hM3Dq, tamoxifen administration was done at P30 and P32.

### Ablation and activation of ChCs

To ablate ChCs for histochemical studies, the expression of DTR in ChCs was induced by intraperitoneally injecting TMX into *Unc5b-CreER:Nkx2.1-Flp:Ai65:Tau-ds-DTR* mice at P21 and P23. Then, diphtheria toxin (DT, 50 μg/kg; Sigma-Aldrich, MO, USA) was intraperitoneally injected at P37 and P39. Histochemical experiments were performed 7 days after DT injection. Clozapine N-oxide (CNO, 0.6 mg/kg; Sigma-Aldrich, MO, USA) was intraperitoneally injected twice a day for 7 days from P46 to P53 to chronically activate ChCs in *Unc5b-CreER* mice infected with AAV-hM3Dq.

### Stereotaxic surgery

Standard surgical procedures were used for stereotaxic injection, as previously described (*27*). Briefly, animals were anesthetized with isoflurane inhalation (1.5% in a mixture with oxygen, applied at 1.0 L/min) and head-fixed in the stereotaxic apparatus. Viruses were injected into dorsal medial prefrontal cortex (dmPFC) at 4 weeks of age, using the coordinates referenced to the Paxinos and Watson Mouse Brain in Stereotaxic Coordinates, 3rd edition: anteroposterior (AP): 1.94 mm; mediolateral (ML): ± 0.25 mm; dorsoventral (DV): –1.7 mm. For each injection site, a total volume of 200 nL of virus was injected at ∼100 nL/min, controlled by hand. The following adeno-associated viral vectors were used to express genes of interest in a Cre-recombinase-dependent manner: AAV2/9-hEF1a-DIO-mCherry-P2A-Tettox-WRPE-pA (6.13 × 10^11^ vg/ml; Taitool, Shanghai, China); pAAV-hSyn-DIO-hM3Dq-EGFP (8.45 × 10^11^ vg/ml; Genechem, Shanghai, China); AAV2/9-hSyn-FLEX-EGFP-WPRE-pA (3.28 × 10^11^ vg/ml; Taitool, Shanghai, China); AAV2/9-hSyn-FLEX-Tdtomato-WPRE-pA (9.85×10^10^ vg/ml, Taitool, Shanghai, China). For immunohistochemical experiments, to minimize the impact of batch effects, AAV virus expressing fluorescent proteins (control) and the ones expressing Tettox or hM3Dq were injected unilaterally in the same mouse but on different sides. For behavioral tests, bilateral injection was applied to achieve robust delivery in ChCs in the mPFC.

### Immunohistochemistry

Animals were anesthetized with isoflurane inhalation (3%) and perfused transcardially with PBS and 4% paraformaldehyde (PFA) in 0.1 mol/L phosphate buffer. The brains were post fixed in 4% PFA in 0.1 mol/L phosphate buffer overnight at 4 ℃ and then dehydrated with 30% sucrose in PBS. Coronal sections at a thickness of 50 μm were prepared using a vibratome (Leica, VT1000S, Wetzlar, Germany). Before incubation with the primary antibody, the brain sections were subjected to antigen retrieval by water bath at 85℃ for 15 min in retrieval buffer (25 mmol/L Tris-HCl, pH 8.5; 1 mmol/L EDTA, 0.05% SDS). After cooling to RT, the sections were washed in PBS for 10 min. Immunohistochemistry was performed as previously described (*71*). Briefly, the sections were blocked with 10% newborn goat serum (NGS) and 0.3% Triton X-100 in PBS at RT for 1 h, followed by incubation in primary antibodies diluted in 3% NGS and 0.3% Triton X-100 in PBS overnight at 4 ℃. The following primary antibodies were used: rabbit anti-RFP (1:10000; Rockland, PA, USA); chicken IgY anti-GFP (1:10000; Aves Labs.Inc, CA, USA); mouse IgG1 anti-Pan Na_v_ 1 (1:100; N419/78; Millipore, MA, USA), mouse IgG1 anti-Na_v_ 1.6 (1:100; K87A/10; NeuroMab, CA, USA); mouse IgG1 anti-gephyrin (1:500; mAb7a; Synaptic Systems, Coventry, UK); guinea pig anti-VGAT (1:500; Synaptic Systems, Coventry, UK); mouse IgG2a anti-AnkyrinG (1:500; N106/36; Millipore, MA, USA); mouse IgG1 anti-parvalbumin (1:500; PVALB-19; (Sigma-Aldrich, MO, USA). After three washes with PBS for 10 min each, the secondary antibodies were applied for 2 h at RT. The secondary antibodies used were (all from Thermo Fisher Scientific, MA, USA): Alexa Fluor (AF) 488 goat anti-mouse IgG1 (1:1000), AF 488 goat anti-mouse IgG2a (1:1000), AF 488 goat anti-guinea pig (1:1000), DyLight 488 goat anti-chicken, AF 555 goat anti-rabbit (1:1000), AF 555 goat anti-mouse IgG1 (1:1000), AF 555 goat anti-mouse IgG2a (1:1000), AF 647 goat anti-mouse IgG2a (1:1000), and AF 647 goat anti-mouse IgG1 (1:1000). The same wash steps were applied after incubation. The brain slices were then mounted onto microscope slides for image acquisition.

### Image acquisition and analysis

Images were acquired from Nikon AX confocal laser scanning microscope (Nikon, Tokyo, Japan) with 10× air-immersion or 60× oil-immersion objective. Images for immunochemical analysis were captured at a resolution of 2048 × 2048 pixels using a Z-series to cover the entire AIS with a depth interval of 0.55 mm. The analysis of AIS was used and modified as previously described (*11, 51*). In brief, the fluorescence profile of AISs were obtained from Fiji (https://fiji.sc/). For each intact AIS, a 3-pixel-wide line was drawn through and past the fluorescence signal of Ankyrin-G in the 60× stacked images when a custom-made Fiji marco (https://github.com/Mojackhak/AIS) was recording the pixel intensity of all z-series slices along the line. The data were import to the MatLab to reconstruct the AIS-containing plane including the AIS profile in the z-direction and gain the 3D profile by the modified published code (*11*). The proximal and distal position of the AIS was identified through the normalized fluorescence intensity of Ankyrin-G dropped below 0.33. The distance of AIS from soma was calculated as the distance between the soma boundary and the proximal position of the AIS. The width of the AIS was recorded at the proximal position of the AIS. The intensity normalization formula is as follows: normalized intensity of the pixel = (pixel intensity − minimum intensity) / (maximum intensity − minimum intensity). For analysis of VGAT and Gephyrin, puncta were manually counted following each single z-plane image of the entire AIS by using Fiji.

### Slice preparation and electrophysiological recordings

The slice preparation and electrophysiological recordings were conducted following previously described methods (*51*). Briefly, mice were anesthetized using isoflurane before decapitation. The mouse brain was rapidly removed and transferred to ice-cold, oxygenated artificial cerebrospinal fluid (section ACSF) (in mmol/L: 110 choline chloride, 2.5 KCl, 4 MgSO4, 1 CaCl2, 1.25 NaH2PO4, 26 NaHCO3, 11 D-glucose, 10 sodium ascorbate, 3.1 sodium pyruvate, pH 7.35, 300 mOsm) for 1 min. Coronal prefrontal cortical slices, 250 µm–300 µm thick, containing the medial prefrontal cortex (mPFC), were cut at 1 °C–2 °C using a vibratome (Leica VT1200S, Wetzlar, Germany). Subsequently, the slices were allowed to recover and were incubated in oxygenated ACSF (in mmol/L: 124 NaCl, 2.5 KCl, 2 MgSO4, 2 CaCl2, 1.25 NaH2PO4, 26 NaHCO3, 11 d-glucose, pH 7.35, 300 mOsm) at 34°C for 30 minutes, and then stayed in room temperature before the slice was transferred into the recording chamber. Whole-cell patch recordings were performed in layer 2/3 of the mPFC, with the subcortical white matter and the corpus callosum used as visual landmarks based on the atlas (Paxinos and Watson, Mouse Brain in Stereotaxic Coordinates, 3rd edition). The patch pipette recording solution contained the following components (in mmol/L): 147 potassium gluconate, 3 KCl, 10 sodium phosphocreatine, 10 HEPES, 4 ATP·Mg, 0.3 Na2GTP, and 0.3 EGTA. The pH of the solution was adjusted to 7.25 with KOH, and the osmolarity was maintained between 290 mOsm and 300 mOsm. Whole-cell patch recordings from pyramidal neurons (PyNs) in layer 2/3 of the medial prefrontal cortex (mPFC) were performed using Axopatch 700B amplifiers (Molecular Devices, Union City, CA). Slices were visualized using an Olympus Bx51 microscope equipped with infrared-differential interference contrast optics (IR-DIC) and a fluorescence excitation source. Throughout the recordings, the chamber was continuously perfused with oxygenated working ACSF at a temperature range of 33°C to 34°C. During voltage-clamp mode, the membrane potential was held at –75 mV, while in current-clamp mode, zero holding current was applied without correcting for the junction potential. The recorded signals were low-pass filtered at 3 kHz, digitized at 80 kHz using a Digidata 1550B digitizer (Molecular Devices), and subsequently analyzed using pClamp 10.7 software (Molecular Devices) and custom-made MATLAB code. To evoke action potentials (APs), a current ramp was employed, and the analysis focused on the first evoked AP. The threshold of an AP was determined as the voltage at which the derivative of the membrane potential (dV/dt) reached 25 mV/ms and was calculated using MATLAB’s spline interpolation function (*51, 72*).

### Behavior assays

The behavior assays were conducted between 10:00 am and 5:00 pm. All subject mice were male and randomly grouped. To minimize stress, all mice were handled for 5 min on 5 consecutive days prior to test day to acclimation to moderate handling. On the test day, the mice were placed in the test room to acclimate to the environment for 30 minutes before the test. At the end of each test, the testing apparatus was scrubbed twice with 75% alcohol and ddH_2_O. Mouse trajectories were recorded and analyzed using Toxtrac (https://toxtrac.sourceforge.io). Acute pharmacogenetic activation of ChCs during the behavior assay was achieved by i.p. injection of CNO (1mg/kg) approximately 30 minutes before the test, while chronic activation of ChCs was achieved by i.p. injection twice a day for 8 consecutive days.

### Social interaction assays

The social testing apparatus comprises a three-chamber Perspex box, with each chamber measuring 60 cm in length, 40 cm in width, and 44 cm in height. The two adjacent chambers are separated by a freely removable baffle. On the day of testing, the subject mouse was placed in the central chamber and allowed to freely explore the empty apparatus for 10 minutes. Empty social boxes were then placed on either side of the apparatus, and the subject mice were allowed to explore them for 10 minutes. In the social ability test, a strange mouse of the same sex and age was randomly placed in one social box, and an object was placed in the social box on the other side. The subject mouse was then allowed to investigate freely for 10 min. In the social novelty test, the object in the social box was replaced with another strange mouse, followed by a 10 min investigation by the subject mouse. We defined a circular area within 5 cm around the social box as the interaction zone and considered the time spent around the interaction zone as interaction time. The degree of preference of a stranger mouse over the object represented the subject’s social ability. The social index was calculated using the following formula: (interaction time with stranger – interaction time with object) / total test time. For the social novelty test, the formula was modified as follows: (interaction time with new mice – interaction time with old mice) / total test time.

### Open-field test

The mice were placed in a plexiglass box measuring 42 cm ×42 cm ×50 cm for 10 minutes to allow for free exploration. Motor performance was quantified by calculating the average speed and total distance traveled. The time spent in the center of the box (half of the box area) was also recorded.

### Elevated plus maze (EPM) test

The EPM apparatus was elevated 73 cm above the floor and consisted of a 6 cm × 6 cm center section, two open arms (35 cm × 6 cm), and two closed arms (35 cm × 6 cm × 19 cm). To assess anxiety-related behavior, the subject mouse was placed in the center section with its head on one side of the open arms, followed by 5 minutes of free exploration. The time spent on the open arms was then recorded.

### Statistical analysis

Statistical analyses and data plots were performed using GraphPad Prism 9 software (GraphPad). All data are presented as mean ±SEM. Unpaired two-tail Student’s *t*-tests were used to compare two groups. Two-way ANOVA was used to compare AP firing frequency and normalized mean fluorescence intensity along AISs. Unpaired multiple *t* test with Holm-Sidak method was used to compare the time course of Tettox and social index. Statistical significance was defined as \**P* <0.05, \*\**P* <0.01, \*\*\**P* <0.001, or \*\*\*\**P* <0.0001. Values of *P* ≥0.05 were considered not significantly different. All figures were composed using Illustrator software (Adobe Systems, USA).

## Acknowledgments

We thank Dr. Martyn Goulding at the Salk Institute for the *Tau-DTR* mice and the Allen Brain Institute for the *Ai65* mice.

## Funding

National Natural Science Foundation of China grant 82071450 and 32371006(Y.T.)

National Natural Science Foundation of China grant 31970971 (M.H.)

National Natural Science Foundation of China grant 32371073 (J.L.)

National Science and Technology Innovation 2030 Major Projects of China STI2030-Major Projects-2022ZD0206500 (M.H.)

STI2030-Major Projects 2021ZD0202500 (Y.S.)

National Natural Science Foundation of China 32130044 and T2241002 (Y.S.)

National Natural Science Foundation of China 32200951 (Y.X.)

China Postdoctoral Science Foundation 2022M720801 (Y.X.)

US NIH NIH R01 MH094705-05 (Z.J.H.)

## Author contributions

Conceptualization: JL, YT

Methodology: RZ, BR, YX, AH, YQ, ZJH.

Investigation: RZ, BR, YX, JT, YZ

Visualization: RZ, JT, YZ, YQ, XX, YT

Supervision: YS, MH, JL, YT

Writing—original draft: RZ, YT

Writing—Review and editing: ZJH, YS, MH, JL, YT

## Data and materials availability

All data are available in the main text or the supplementary materials.

