## supplementary figures for "Axo-axonic synaptic input drives homeostatic plasticity by tuning the axon initial segment structurally and functionally"

**Fig. S1.**

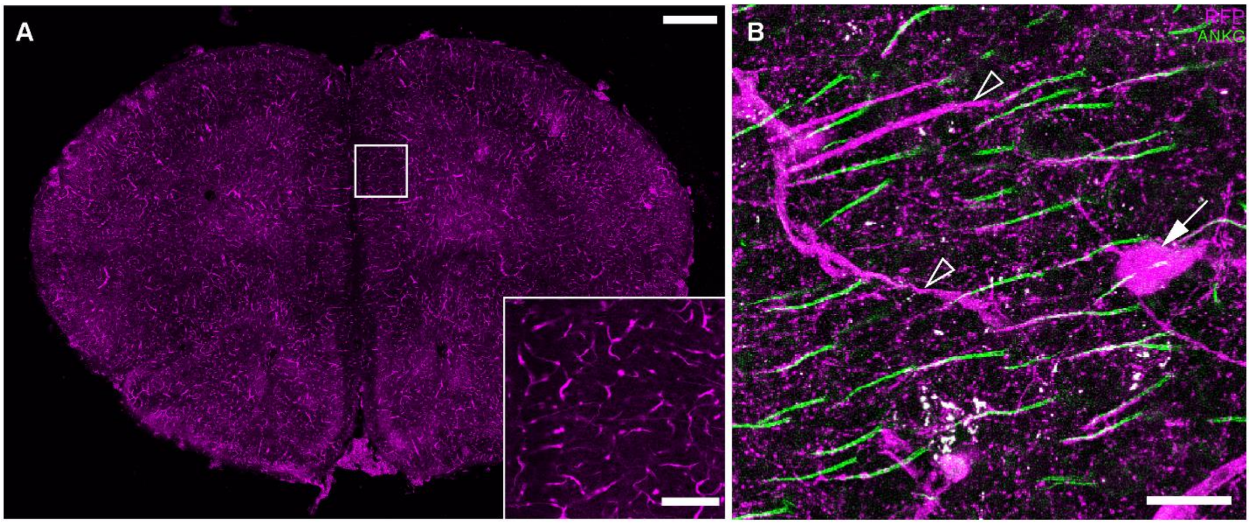


Supplementary Figure 1

**The *Unc5b-CreER:Ai14* mouse line labels ChCs and capillary endothelial cells.**
**(A)** Representative image of the *Unc5b-CreER:Ai14*-labeled forebrain slice. Scale bar: 500 µm (upper); 100 µm (lower). **(B)** Immunostaining against RFP and AnkG in the L2/3 mPFC of *Unc5b-CreER:Ai14* mice. Scale bar: 20 µm. The arrow indicates the ChC cell body, and arrowhead indicates capillary endothelial cells.

**Fig. S2.**

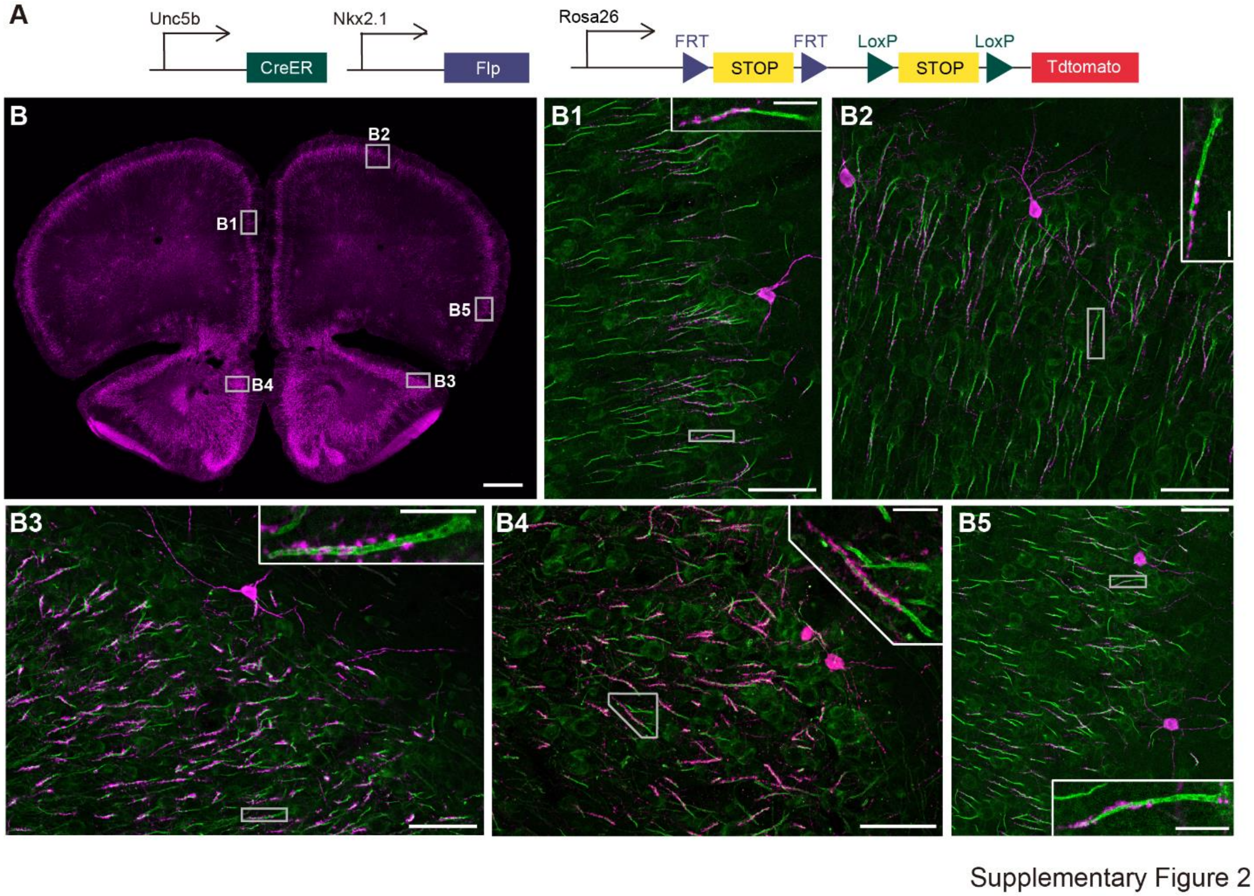


**The *Unc5b-CreER:Nkx2.1-Flp:Ai65* mouse line labels ChCs with high efficiency.**
**(A)** Schematic showing the strategy of intersectional labeling (tdTomato) of the ChCs. **(B)** Representative images of immunostaining against RFP (ChCs) and AnkG (AISs) in the prelimbic cortex (B1), motor cortex (B2), piriform cortex (B3 and B4), and agranular insular cortex (B5) in *Unc5b-CreER:Nkx2.1-Flp:Ai65* mice. Scale bar: 50 µm in B1-5; 10 µm in insets of B1-5.

**Fig. S3.**

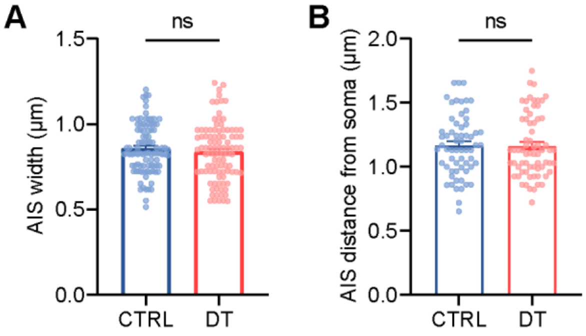


Supplementary Figure 3

**No changes in AIS width and the distance from soma after ChCs ablation.**

**(A and B)** No significant changes in AIS width **(A)** and distance from soma **(B)** between control and DT-treated group (In **(A)**, CTRL: $n$ = 95 cells, 3 mice; DT: $n$ = 97 cells, 3 mice, $P$ = 0.369, unpaired $t$-test; In **(B)**, CTRL: $n$ = 62 cells, 3 mice; DT: $n$ = 67 cells, 3 mice, $P$ = 0.896, unpaired $t$-test). Data are represented as mean ± SEM. n.s. $P>0.05$.

**Fig. S4.**

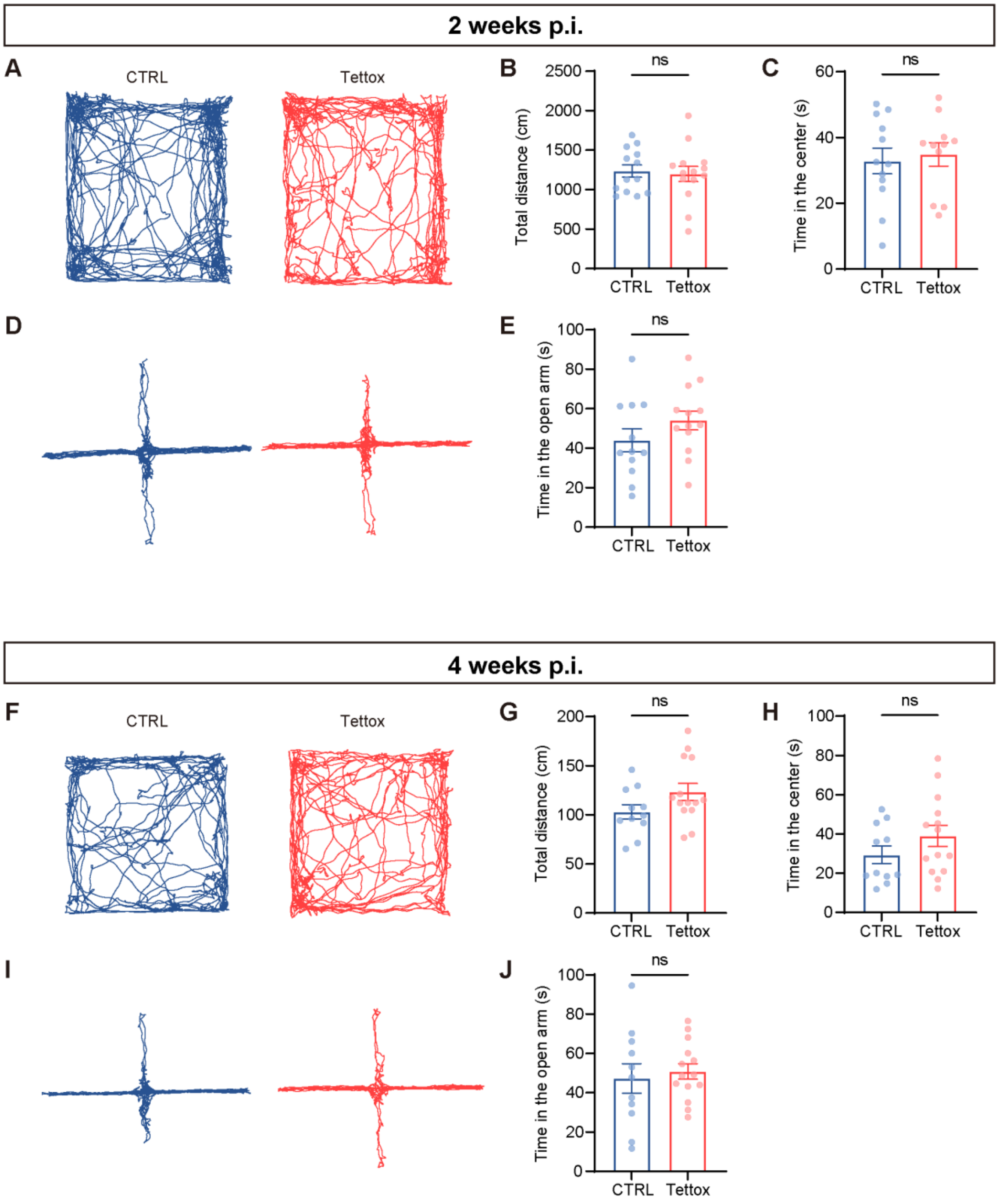


Supplementary Figure 4

**No detectable anxiety-like behaviors after blocking ChCs synaptic transmission.** Representative trajectory of open field test 2 weeks (acute) after the animals were injected with AAV-DIO-tdTomoato (CTRL) or AAV-DIO-Tettox. **(B and C)** No changes in the total moving distance **(B)** or the time spent in the center area **(C)** in the indicated groups. **(D)** Representative trajectory of elevated plus maze 2 weeks (acute) after animals were injected with AAV-DIO-tdTomoato (CTRL) or AAV-DIO-Tettox. **(E)** No difference in the time spent in the open arm in the indicated groups. **(F)** Representative trajectory of open field test 4 weeks (chronic) after the animals were injected with AAV-DIO-tdTomoato (CTRL) or AAV-DIO-Tettox. **(G and H)** No changes in the total moving distance **(G)** or the time spent in the center area **(H)** in the indicated groups. **(I)** Representative trajectory of elevated plus maze 4 weeks (chronic) after the animals were injected with AAV-DIO-tdTomoato (CTRL) or AAV-DIO-Tettox. **(J)** No difference in the time spent in the open arm in the indicated groups. $n$ = 12 for CTRL and 11 for Tettox in the acute condition; $n$ = 11 for CTRL and 14 for Tettox in the chronic condition. unpaired *t*-test, *n.s.* $P>0.05$. Data are represented as mean ± SEM.

**Fig. S5.**

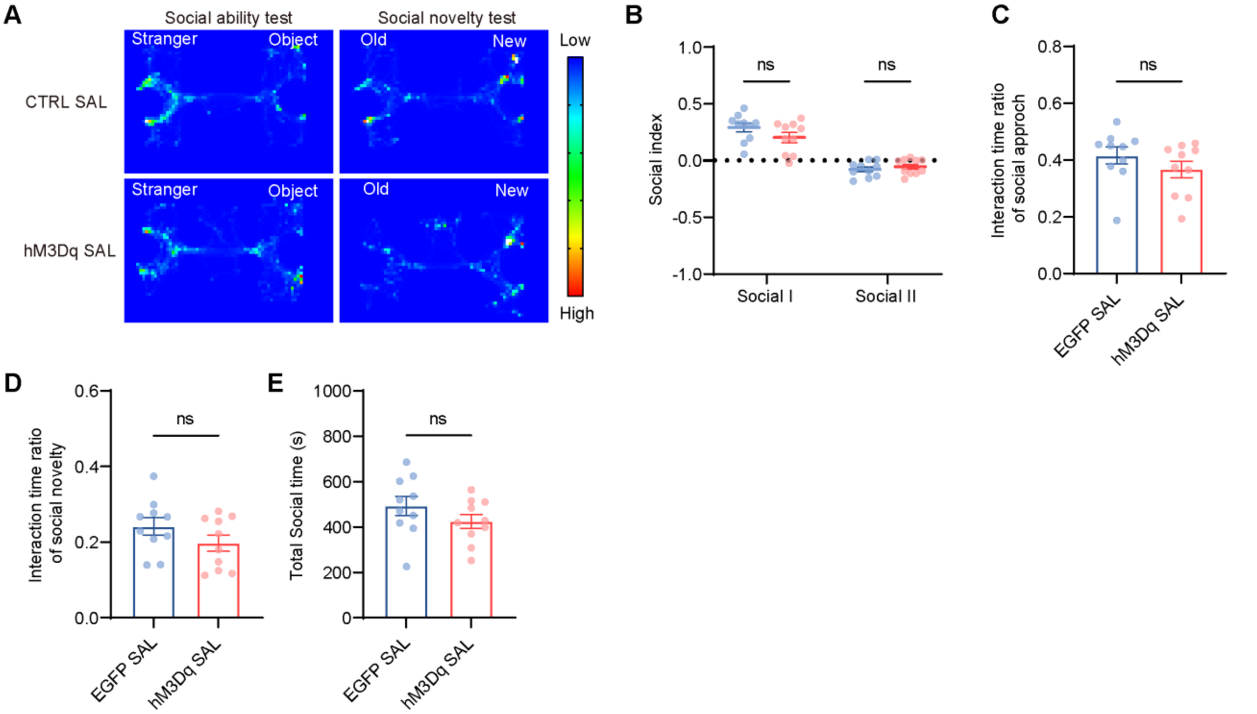


Supplementary Figure 5

**Social performance was not changed by expression od hM3Dq without CNO activation.** **(A)** Representative heatmaps of social ability and social novelty test in animals expressing AAV-DIO-tdTomato and AAV-DIO-hM3Dq without CNO activation. （SAL: saline） **(B)** Unchanged preference for animals in social approach and social novelty in the SAL group (*P* = 0.276 for social approach and 0.421 for social novelty, unpaired Holm-Sidak multiple *t*-test). **(C and D)** The interaction time of social approach and social novelty were not changed in the indicated groups (*P*>0.05, unpaired t test). **(E)** No significant difference in the total social time in the SAL group (*P* = 0.208, unpaired t test). Data are represented as mean ± SEM.
